# Peptidyl Protease Inhibitors Block ERM-BP Function and Suppress *Entamoeba* Encystation

**DOI:** 10.64898/2026.09.16.752027

**Authors:** Shreyasee Hazra, Suman Kalyan Dinda, Somashree Pandit, Dipak Manna

**Affiliations:** Department of Biomedical Science and Technology, School of Biological Sciences, Ramakrishna Mission Vivekananda Educational and Research Institute (RKMVERI), Kolkata, India

**Keywords:** Amoebiasis, Encystation, Peptidyl Protease, Transcription Factor

## Abstract

*Entamoeba* encystation process is a crucial developmental process that support to parasite persistence and disease transmission, yet the molecular mechanisms governing this stage conversion remain poorly understood and represent an attractive opportunity for therapeutic intervention. The Encystation Regulatory Motif-Binding Protein (ERM-BP), a transcription factor and key regulator of encystation, therefore represents a promising target for anti-amoebic drug development to prevent disease transmission. In the present study, we applied a target-based drug discovery approach to screen small molecules that can block the activity of ERM-BP, with a particular focus on the Cys-198 residue of this protein, which is functionally important. A library comprising 100 compounds including broad-spectrum nicotinamidase inhibitors and cysteine-reactive electrophiles were screened against ERM-BP by Protein thermal shift assays (PTSA). Thirty-six compounds bound to wild-type ERM-BP while analysis using the C198A mutant narrowed the candidates to three compounds exhibiting Cys-198-dependent interactions. These three peptidyl protease inhibitors, PFMK (Z-Ala-Phe-FMK), LEK (Z-Leu-EK), and EBLL (Ethylbenzyl-Leu-Lys) binds to ERM-BP-WT and abolishes the DNA-binding activity of ERM-BP in a concentration-dependent way. Cellular evaluation further revealed distinct activity of PFMK, LEK and EBLL on trophozoite growth and PFMK and EBLL reduced encystation efficiency significantly, produced irregular, and structurally defective cysts, which fails to excyst to trophozoites, whereas LEK had no significant effects on encystation efficiency or cyst morphology. Altogether, these findings identify peptidyl protease inhibitors PFMK and EBLL as promising compounds to target ERM-BP and block the *Entamoeba* encystation, providing a framework for developing therapeutics targeting parasite differentiation and encystation.

## INTRODUCTION

Amoebiasis, caused mainly by the protozoan parasite *Entamoeba histolytica*, is still a major global health issue, especially in areas with poor sanitation and limited access to clean water (1–3). The parasite has two developmental forms: the motile trophozoite, which is responsible for colonizing the body and invading tissues, and the environmentally resistant cyst, which enables transmission from one host to another (4, 5). *Entamoeba* transmitted to human through fecal-oral route by the ingestion of cyst with contaminated food and water (6–8). After being ingested, the cysts undergo excystation in the host’s intestine to release trophozoites, which can then colonize the intestinal tract and, in certain instances, lead to invasive disease. Encystation is therefore essential both for the survival of the parasite outside the host and for the spread and sustaining of the infection cycle (9–12). If encystation could be prevented, parasite transmission would be interrupted and environmental contamination would be reduced as well, making it a promising therapeutic approach (13–15).). Even though encystation is of great biological and epidemiological significance, the molecular mechanisms that control it in *Entamoeba* have not yet been fully understood. The process of encystation includes extensive cellular and transcriptional reprogramming, such as alterations in gene expression, metabolism, cell shape, and the production of components that are specific to the cyst (16–20). Transcription factors that coordinate these developmental changes are probably playing a central role in the regulation of cyst formation (21–23). If such regulatory proteins could be identified and their functions determined, this might open up new possibilities to understand parasite differentiation and for developing interventions that specifically interfere with transmission (24–26).

The Encystation Regulatory Motif-Binding Protein (ERM-BP) is a transcription factor which has been involved in the regulation of encystation in *Entamoeba* and thus may be an important part of the transcriptional system responsible for this developmental change (9, 27). By controlling the expression of the genes associated with encystation, ERM-BP play a crucial role in the coordinated molecular processes needed for cyst formation. The mutation studies identified a functionally important cysteine residue in the nicotinamidase domain of ERM-BP at Cys-198 position, that is crucial for its function. A single amino acid mutation at the Cysteine residue of ERM-BP at position 198 to Alanine (C198A) impairs both NAD^+^ and DNA binding activity. It further leads to protein mis-localization in both trophozoites and cysts and significantly reduce encystation efficiency (28). So, targeting ERM-BP using broad spectrum nicotinamidase inhibitors and cysteine-reactive electrophiles could have huge potential blocking the encystation of *Entamoeba* which can eventually stop the transmission of the disease to the next host. Since ERM-BP is a developmental regulator specific to the parasite, it is a notably promising therapeutic target because it could offer high selectivity. Target-based drug discovery delivers a structured method for finding compounds that directly interact with and modify specific parasite proteins. In order to be developed as a targeted therapy, both its molecular target and functional validation are required and for an example in a recent target-based drug discovery approach fumagillin demonstrates its anti-amoebic activity by means of covalent inhibition of EhMetAP2 (29). Compound screening can be used to identify potent anti-*Entamoeba* molecules against drug-resistant parasites implying high-throughput techniques (HHT) (30–32). One such HHT is Protein thermal shift assays (PTSA) which is widely used to detect changes in protein stability caused by ligands and thus aid in the screening of compound libraries for evidence of target engagement (33, 34). Significantly, comparing how compounds interact with wild-type and site-directed mutant proteins can provide evidence for binding at a specific residue and assist in identifying the key molecular features that determine ligand recognition. In the case of ERM-BP, compounds that are reactive towards cysteine can be used to alter the function of transcription factor ERM-BP. In addition to its function as a transcriptional regulator, ERM-BP has a number of structural characteristics that make it suitable for rational drug design. Biochemical and structural analyses have identified a highly conserved cysteine residue (Cys-198) that plays a critical role in maintaining the protein’s functional integrity (27, 28). The nucleophilic thiol-side-chain of Cysteine provides an attractive site for selective covalent modification by electrophilic small molecules. Covalent inhibition has been reported in recent drug discovery approaches because it enables prolonged target occupancy, improved pharmacodynamic properties, and exceptional target selectivity when directed toward uniquely positioned reactive residues (35, 36). Several successful clinical drugs including inhibitors of kinases, proteases, and other enzymes exploit this principle by targeting reactive cysteine residues within their active or regulatory sites. Despite these advances, the application of cysteine-directed covalent chemistry to protozoan transcriptional regulators remains largely unexplored (37, 38). The growing success of covalent drug discovery has been driven by the development of carefully optimized electrophilic warheads that selectively modifying target cysteine residues while minimizing nonspecific protein alkylation (39). Functional groups such as fluoromethyl ketones, epoxides, acrylamides, vinyl sulfones, and related electrophiles have demonstrated remarkable efficacy against diverse biological targets (40–42). The fact that reversible molecular recognition is followed by irreversible engagement of the target gives rise to both potency and specificity, which is why electrophilic compounds are especially attractive as candidates for targeting ERM-BP (43).

In this study we used a target-directed small-molecule screening method in order to find compounds which can interact with ERM-BP, focusing especially on Cys-198. A library of 100 compounds comprising broad-spectrum nicotinamidase inhibitors and cysteine-reactive electrophiles was screened against wild-type ERM-BP using PTSA, followed by secondary screening with the ERM-BP-Mutant (C198A) to identify Cys-198-dependent interactions (**Supplemental Table 1**). Candidate compounds were subsequently evaluated for their effects on ERM-BP DNA-binding activity, trophozoite growth, and encystation. This integrated approach led to the identification of PFMK and EBLL as lead compounds that interfere with encystation and induce morphological abnormalities in developing cysts. PFMK (Z-Ala-Phe-FMK), LEK (Z-Leu-EK), and EBLL (Ethylbenzyl-Leu-Lys) are peptidyl protease inhibitors designed to interact with protease substrate-binding sites and, in the case of electrophilic derivatives, reactive catalytic residues such as cysteine. PFMK contains a fluoromethyl ketone (FMK) warhead capable of covalent modification of catalytic cysteines, whereas LEK and EBLL contain peptide-mimetic motifs that may facilitate target recognition. Z-FA-FMK (PFMK) is also found a potent, irreversible inhibitor of a broad range of cysteine proteases, including cathepsins B, L, and S, cruzain, and papain. Previous studies have demonstrated that PFMK suppresses the production of pro-inflammatory cytokines, including IL-1α, IL-1β, and TNF-α, in LPS-stimulated macrophages, primarily through inhibition of NF-κB signaling (44–46). PFMK possesses significant antiviral activity against SARS-CoV-2 (47). Moreover, PFMK has been reported to exhibit potent inhibitory activity against specific strains of mammalian reovirus, further highlighting its potential as a broad-spectrum antiviral compound (48). Interestingly, our findings PFMK as ERM-BP-interacting compounds suggests that these scaffolds may also engage non-proteolytic targets. These findings provide evidence for the chemical tractability of ERM-BP and establish a framework for targeting parasite-specific transcriptional regulators to disrupt *Entamoeba* differentiation and transmission. By focusing on inhibiting cyst formation rather than solely eliminating trophozoites, this study provides a conceptual framework for the development of next-generation anti-amoebic therapeutics that block parasite transmission, reduce environmental persistence, and ultimately contribute to more effective control of amoebiasis.

## RESULTS

### Target-Based Screening of Small Molecules Against ERM-BP

We employed target-based drug discovery approaches directly targeting the encystation specific Transcription Factor ERM-BP (Encystation Regulatory Motif-Binding Protein), with particular emphasis on compounds targeting its functionally important cysteine residue (Cys-198). 100 compounds including broad spectrum nicotinamidase inhibitors and a variety of cysteine-reactive electrophiles were screened against wild-type ERM-BP. The workflow of compound screening and follow up validation is summarized in **Fig. 1A**. In the first step 100 compounds were screened against recombinant wild-type ERM-BP (ERM-BP-WT) by the Protein Thermal Stability Assay (PTSA), which measures ligand-induced changes in protein thermal stability. If a small molecule binds to a protein it can result in stabilization or destabilization of the protein structure, producing a measurable shift in its melting temperature (Tm), thereby enabling rapid identification of potential binders. Each compound was evaluated at a final concentration of 100 μM against purified recombinant ERM-BP-WT (**Supplemental Fig. 1**). PTSA analysis identified 36 compounds that produced a significant thermal shift relative to the DMSO treated as control, indicating detectable interaction with ERM-BP. The remaining 64 compounds failed to induce any significant change in protein thermal stability and were therefore excluded from further investigation. Structural analysis of ERM-BP revealed that the protein contains an N-terminal DNA-binding domain and a C-terminal nicotinamidase domain. Within the nicotinamidase domain, cysteine residue C198 was identified as a critical residue involved in NAD⁺ binding as shown in **Supplemental Fig. 2A**. As Cys-198 residue of ERM-BP-WT has previously been identified as a critical residue for its function and represents a potential site for covalent modification, the 36 primary hits were subjected to a secondary screening using a recombinant ERM-BP-Mutant (C198A), in which cysteine-198 was replaced with alanine (15). This C198A mutation abolishes the reactive thiol group while preserving the overall protein structure, thereby allowing discrimination between compounds that bind specifically through Cys-198 and those that interact elsewhere on the protein. Comparison of the PTSA profiles obtained with ERM-BP-WT and ERM-BP-mut revealed that 3 compounds (PFMK, LEK and EBLL) out of 36 compounds no longer exhibited detectable binding to the C198A mutant. The loss of interaction following substitution of Cys-198 residue to Ala strongly suggests that these compounds likely interact directly with the thiol group of Cys-198 through a covalent or cysteine-dependent mechanism. However, the remaining 33 compounds retained binding to both the wild-type ERM-BP-WT and mutant proteins ERM-BP-mut (C198A), indicating that their interaction with ERM-BP is independent of Cys-198 and is likely mediated through alternative binding sites or non-covalent interactions **Fig. 1B**. Altogether, these two-steps target-based screening approach efficiently narrow down an initial library of 100 electrophilic molecules to 3 high-confidence Cys-198-dependent ERM-BP binders. These compounds PFMK, LEK and EBLL were prioritized for subsequent biochemical, biophysical and cellular characterization to evaluate their ability to inhibit ERM-BP function, disrupt encystation-associated signaling, and suppress *Entamoeba* growth and differentiation. As shown in **Fig. 1B** the fluorescence signal and first-derivative melting curves (dFI/dT), the electrophilic inhibitors PFMK (Z-Ala-Phe-FMK), LEK (Z-Leu-EK), and EBLL displayed characteristic interactions with the protein ERM-BP-WT driven by their active functional warheads (fluoromethyl ketone and epoxide groups highlighted in red). The docking scores of the different compounds (PFMK, LEK, EBLL, and NAD^+^) with ERM-BP were evaluated. The predicted docking interaction between PFMK and ERM-BP is shown in Supplemental Fig. 2B and the interacting amino acid residues highlighted in the inset. A prominent rightward shift in the dFI/dT denaturation peaks were observed in the presence of ERM-BP WT compared to the DMSO control, confirming successful ligand-induced thermal stabilization. On the contrary, when counter-screened against the ERM-BP-Mut (C198A), these compounds showed no rightward shift, showed overlapping fluorescence similar to DMSO control, suggesting these compounds does not bind to ERM-BP-WT through C198 (**Fig. 1B)**.

**Figure 1.**
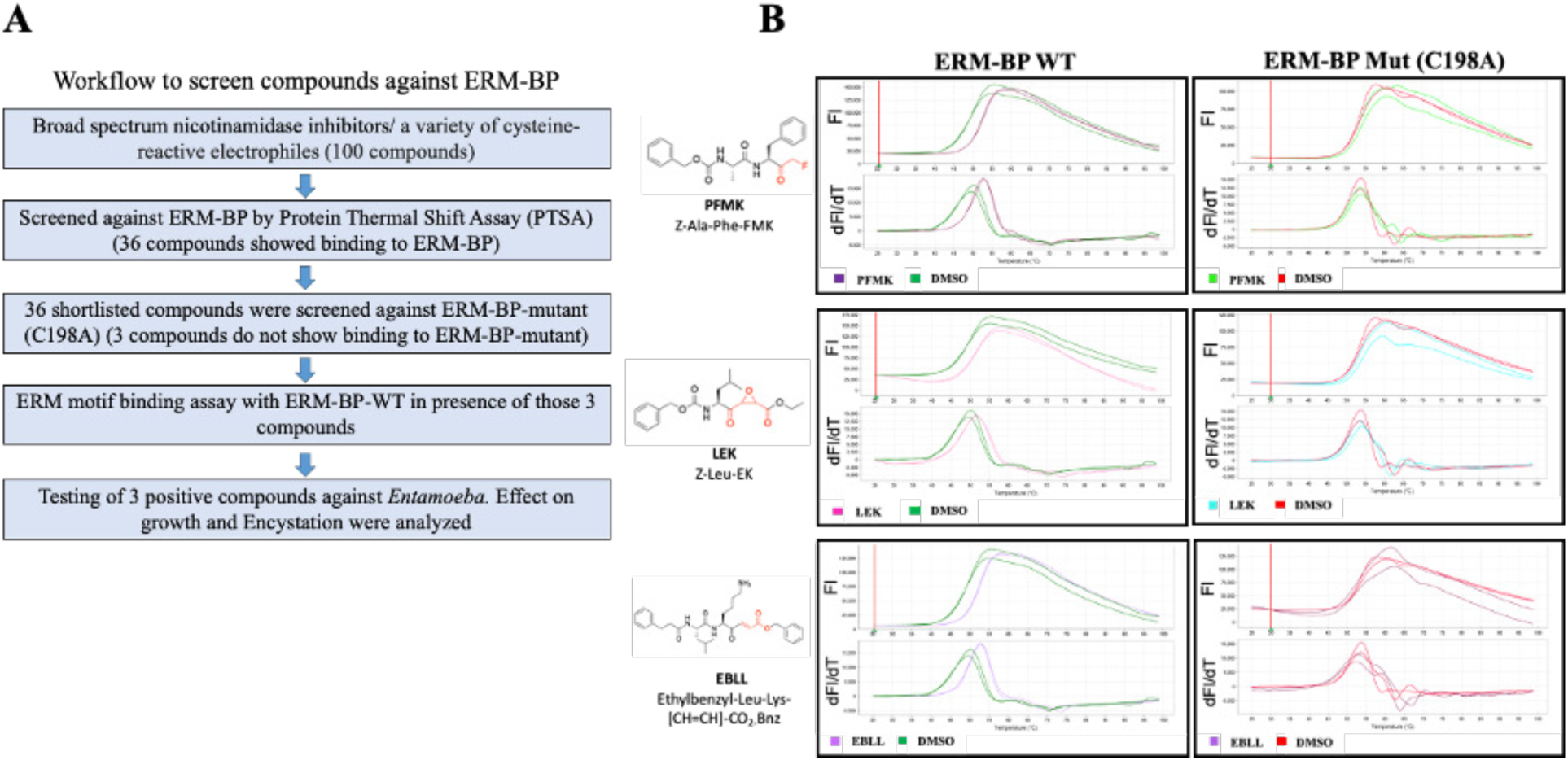
Systematic Screening workflow for the identification of inhibitors targeting ERM-BP and Protein Thermal Shift Assay (PTSA) profiles with the selected compounds. **(A)** The schematic depicts the sequential steps utilized to isolate and validate potent compounds against the *Entamoeba* Encystation Regulatory Motif Binding Protein (ERM-BP). The workflow initiates with a library of 100 compounds consisting of broad-spectrum nicotinamidase inhibitors and cysteine-reactive electrophiles. Primary screening was performed using a Protein Thermal Shift Assay (PTSA) against wild-type ERM-BP. 36 primary hits were identified and these were subjected to a secondary screening using a recombinant ERM-BP mutant (C198A). Three compounds showing Cys-198-dependent interactions and do not show any shift against C198A in PTSA. Downstream verification of these three compounds includes an ERM motif-binding assay using wild-type protein (ERM-BP-WT) in presence of compounds showing selective binding, culminating in functional *in vitro* evaluation via *Entamoeba* viability assays and encystation inhibition studies. **(B)** Thermal denaturation profiles showing raw fluorescence intensity (FI, top panels) and first-derivative melting curves (dFI/dT, bottom panels) as a function of temperature (20 to 100°C). The panel columns compare compound interaction with wild-type ERM-BP (ERM-BP WT, left) versus the cysteine mutant variant ERM-BP-Mut (C198A, right). Rows from top to bottom track the chemical structures and thermal shifts induced by PFMK (Z-Ala-Phe-FMK), LEK (Z-Leu-EK), EBLL. Distinct rightward shifts in the derivative peak temperature (Tm) indicate ligand-induced thermal stabilization, while an absence of a shift between WT and Mutant variants highlights non-specific or independent interaction mechanisms.

### PFMK inhibits DNA-binding activity of ERM-BP in a concentration-dependent manner as determined by Electrophoretic Mobility Shift Assay (EMSA)

To investigate whether these three inhibitors PFMK (Z-Ala-Phe-FMK), LEK (Z-Leu-EK), and EBLL interferes with the nucleic acid-binding activity of the target protein, electrophoretic mobility shift assays (EMSA) were performed using a labeled probe containing the cognate binding sequence. Incubation of the labeled probe with the purified protein generated a prominent retarded band corresponding to the target-protein complex, verifying a highly specific interaction compared to the rapidly migrating free probe. All these three compounds interfere DNA binding property of ERM-BP (**Fig. 2A)**. However, PFMK showed the most significant effect and when evaluating the competitive effects of PFMK across a concentration gradient, the compound progressively reduced the intensity of this shifted complex in a dose-dependent manner (**Fig. 2B)**. This loss of complex formation was accompanied by a corresponding, reciprocal increase in the intensity of the free probe, demonstrating that PFMK successfully displaces the protein from its target sequence. The gradual disappearance of the shifted complex without the formation of secondary smeared or nonspecific bands indicates that the inhibition stems from target-specific blocking rather than protein aggregation or probe degradation. Overall, these findings demonstrate that PFMK effectively suppresses target protein-binding activity in vitro, providing strong mechanistic evidence that it disrupts functional downstream interactions by directly competitive or allosteric inhibition of its binding pocket.

**Figure 2.**
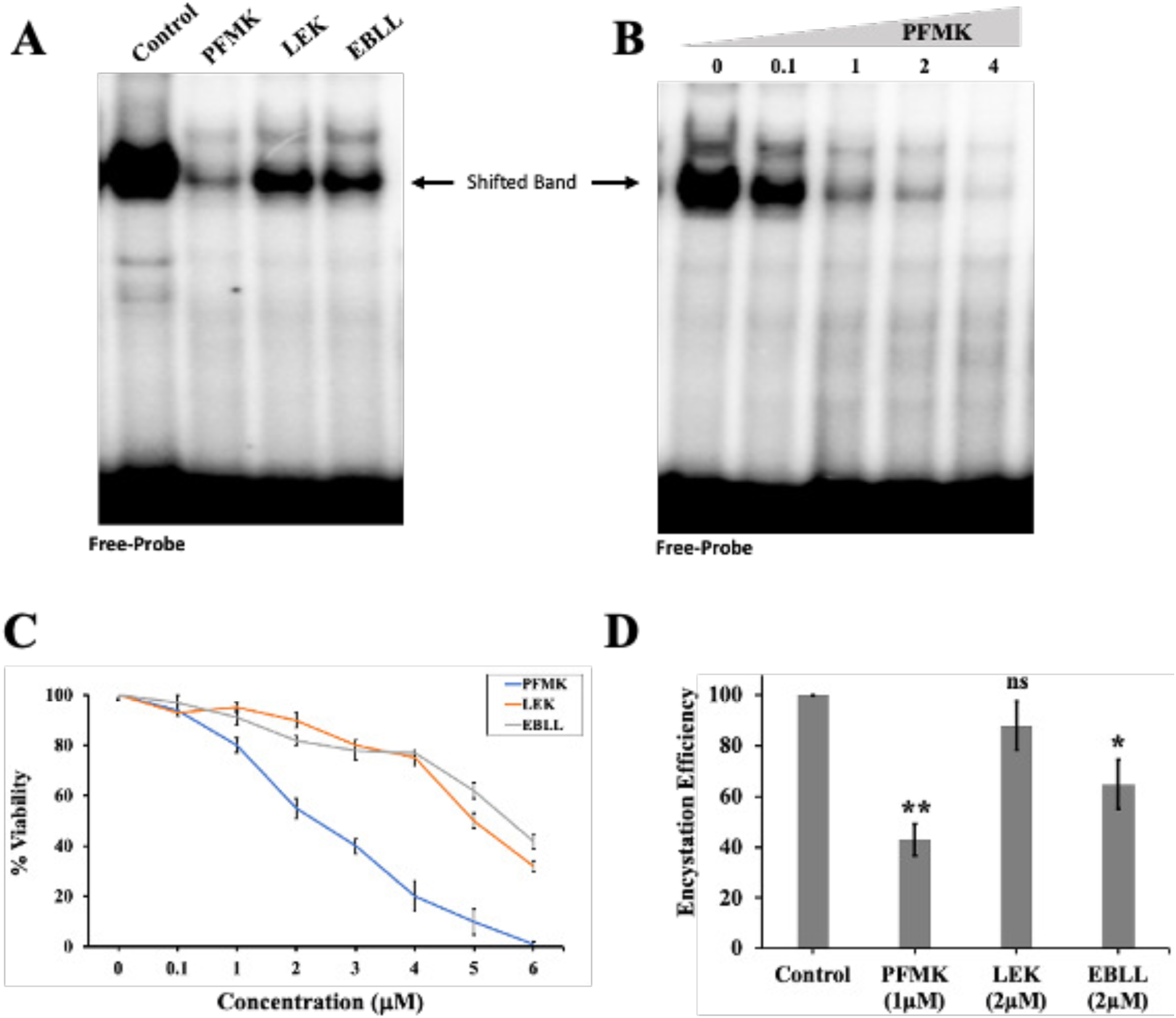
Peptidyl protease inhibitors block DNA-binding activity of ERM-BP and affects *Entamoeba* growth and encystation. **(A)** Representative EMSA showing the DNA-binding specificity of the target protein. The labeled DNA probe was incubated with the purified protein in the absence or presence of appropriate control reactions (negative control, protein alone, and specificity controls such as unlabeled competitor DNA or mutant probe, as indicated above the lanes). Formation of the DNA–protein complex is observed as a shifted band, whereas the lower band corresponds to the free DNA probe. **(B)** Representative EMSA demonstrating the effect of increasing concentrations (0 to 4 μM) of PFMK on DNA-binding activity. The DNA-binding reaction was performed in the presence of increasing concentrations of PFMK (indicated above the lanes). A progressive reduction in the intensity of the shifted DNA–protein complex was observed with increasing compound concentration, accompanied by a corresponding increase in the free probe, indicating concentration-dependent inhibition of protein–DNA interaction. The blot shown is representative of at least three independent experiments. **(C)** *Entamoeba* cell viability was measures across a range of concentrations (0 to 6 μM) for three compounds: PFMK (blue), LEK (orange), and EBLL (grey). Error bars represent the standard deviation of the mean (SEM). **(D)** Encystation efficiency was measured following treatment with a control (DMSO), 1 μM PFMK, 2 μM LEK, and 2 μM EBLL. Data are mean ±s.d. (n = 3) Student’s t-test; (∗p<0.05, ∗∗p<0.01), while “ns” denotes no statistically significant difference.

### Drug activity against encystation and mature cysts

To evaluate the effect of these three compounds on Entamoeba trophozoites, a dose-response viability assay (0 to 6 μM) was performed. As shown in **Fig. 2C**, all three tested compounds exhibited loss of cell viability. PFMK (blue) demonstrated a highly pronounced, dose-dependent decrease in cell viability with an IC50 value of 2.5 μM; while viability LEK (orange) and EBLL (grey) comparatively less potent with IC50 value of 5.0 and 5.5 μM. To determine whether these compounds could block encystation sub-lethal concentrations of the compounds were treated (1 μM for PFMK; 2 μM for LEK and EBLL) for 72 h where at least 80%-90% cells are viable, DMSO (1%) treated as solvent control (100%) (**Fig. 2D**). Among these three compounds, PFMK emerged as the most potent inhibitor of encystation and encystation efficiency down significantly to approximately 43% (p<0.01). EBLL (2 μM) demonstrated a moderate yet statistically significant efficacy (p<0.05), successfully reducing encystation efficiency to roughly 65%. Whereas, LEK (2 μM) showed nominal inhibitory action, dropping encystation efficiency only slightly to ∼88%, an effect that failed to achieve statistical significance (ns).

### PFMK and EBLL treated cysts show altered cyst morphology

In the DMSO control and LEK-treated groups, most cells formed normal, mature cysts. These cysts were round, symmetrical, and surrounded by a bright, thick, and uniform blue wall, indicating healthy cyst formation (**Fig. 3A**). DMSO control and LEK cysts contained strong DNA signals typical of mature cysts. In contrast, PFMK and EBLL cysts showed much weaker DNA staining, indicating poor development and immature cyst formation (**Supplemental Fig. 3A**). In contrast, PFMK and EBLL treatment produced fewer cysts, and many appeared abnormal with shattered cell wall. Staining the cyst outermost wall by *Entamoeba* Jessie-3 antibody showed even more severe defects as shown in **Fig. 3B**, including collapsed or wrinkled walls and patchy staining, suggesting major structural damage and cyst wall formation and these cysts are failed to excyst to form trophozoites (**Supplemental Fig. 3B**). Overall, PFMK and EBLL disrupted normal cyst development, producing defective and immature cysts, whereas LEK had little or no inhibitory effect and resembled to DMSO control. The mechanistic function of PFMK and EBLL are shown in working model in **Fig. 4** and depicts how these compounds target ERM-BP, disrupt its function, inhibit cyst-specific gene expression and eventually block the transmission of the disease.

**Figure 3.**
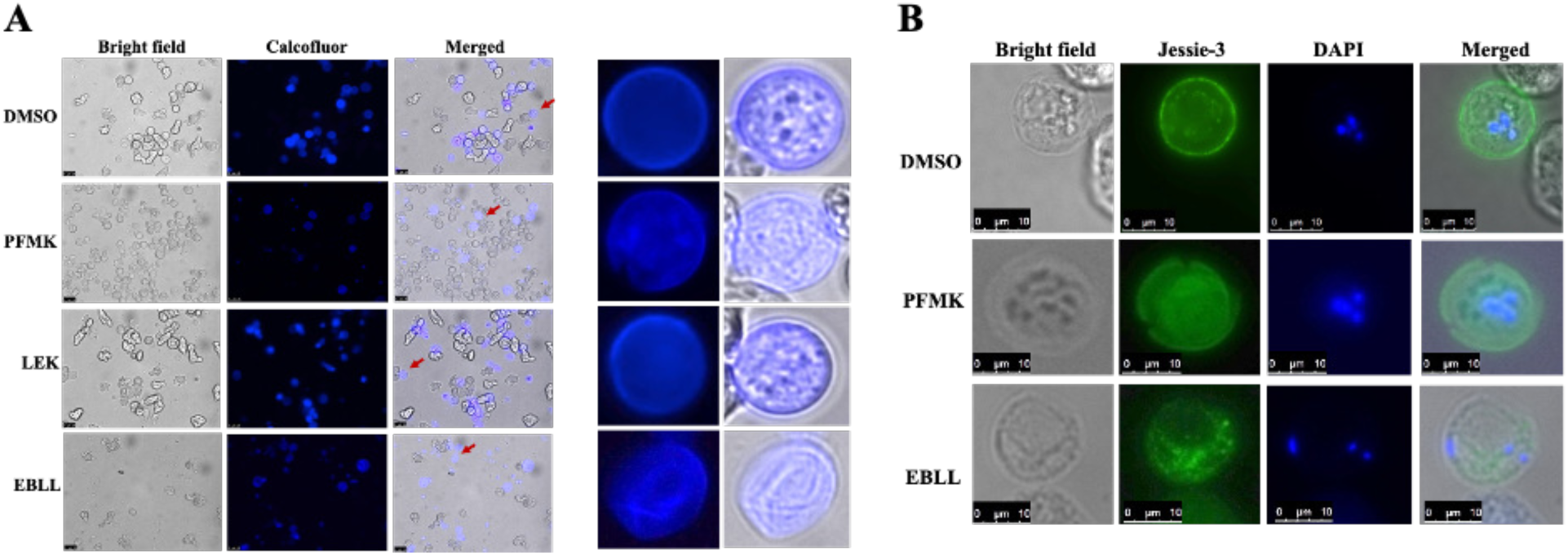
PFMK and EBLL leads to the formation of abnormal and immature cysts. **(A)** *E. invadens* cells were treated with compounds (PFMK and EBLL) or DMSO as Control and encysted for 72 hr, cells were stained with calcofluor white (Blue) and Propidium Iodide (PI) (Red). Mature control cysts exhibit well-defined, robust chitin walls and quadri-nucleated cysts. Cysts treated with PFMK and EBLL display disrupted, faint, or compromised cyst walls and abnormal nucleated cysts, indicative of immature cyst formation. Scale bars 25 μm. **(B)** PFMK and EBLL treated cells show disintegrated localization of Jessie3 (green), nuclear DNA was stained with DAPI (blue). Scale bars represent 10 μm.

**Figure 4.**
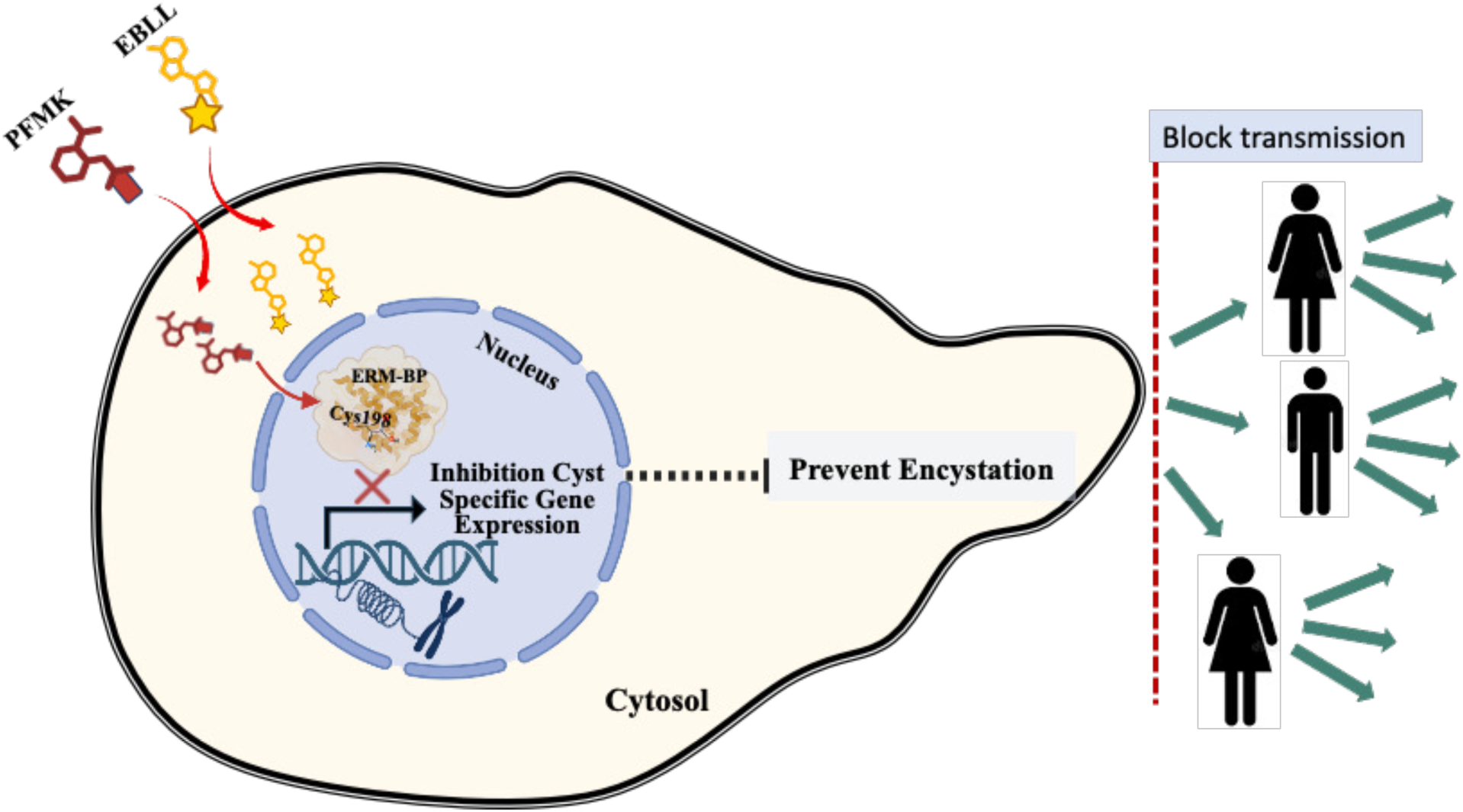
Schematic representation of ERM-BP-inactivation by small molecules that leads to inhibit the cyst-specific gene expression and subsequently encystation. The right panel illustrates the resulting blockade of cyst transmission, thereby potentially limiting the transmission of the disease.

## DISCUSSION

Encystation is an important developmental stage in the biology of *Entamoeba* and plays a key role in the transmission of the disease. Although, this stage conversion from trophozoites to cysts is biologically important, a very few molecular targets associated with encystation have been looked at for the development of therapy. This study concentrates on ERM-BP, a transcription factor which has previously been reported to be involved in the regulation of encystation-specific genes. The use of PTSA made it possible to quickly and efficiently find small molecules that can interact with ERM-BP and inhibit its function. Of the 100 compounds tested, 36 were found to show a detectable interaction with ERM-BP-WT, which means that the interaction is relatively selective. Importantly, secondary screening using the mutant ERM-BP (C198A) allowed the identification of compounds that are likely acting on the key cysteine residue at position 198. This C198A screening exclude three compounds PFMK, LEK, and EBLL that show selective binding with ERM-BP-WT only and no interactions with C198A suggests that the interaction depends on cysteine 198. Since the C198A mutation led to a loss of compound binding, Cys-198 is probably a reactive and functionally important residue in ERM-BP. Covalent targeting of cysteine residues has now become a powerful strategy in drug discovery as a result to its greater specificity and the fact that it leads to a prolonged interaction with the target. Of the compounds identified, PFMK and EBLL stand out as especially promising since they have a strong interaction with ERM-BP-WT and show no binding to the mutant protein. These compounds could interfere with the transcriptional regulation carried out by ERM-BP and therefore inhibit encystation. As cyst formation is an important step for parasite transmission, preventing encystation could greatly reduce the spread of amoebiasis. The high-throughput encystation assay that has been optimized in this study provides an efficient way of evaluating anti-encystation compounds. Collectively, our findings highlight parasite-specific transcriptional regulators as promising therapeutic targets for interfering with encystation, thereby offering a potential strategy to reduce disease transmission of *Entamoeba* and the overall burden of amoebiasis.

## CONCLUSION

The study shows that a target-based approach to drug discovery can be used to find small-molecule inhibitors of the transcription factor ERM-BP in *Entamoeba*. When 100 compounds this group including nicotinamidase inhibitors and cysteine-reactive electrophilic compounds were screened using PTSA, three of the compounds were found to interact with ERM-BP in a manner that is dependent on Cys-198. Of these, PFMK and EBLL displayed the most promising characteristics, showing a selective interaction with wild-type ERM-BP but not with the C198A mutant; this indicates that Cys-198 plays an important role in the binding of the compounds. The results show preliminary evidence that ERM-BP is chemically tractable and may therefore serve as a possible molecular target for interfering with *Entamoeba* encystation. Further validation using *in vitro* encystation assays and *E. histolytica* clinical isolates will be required to establish the biological efficacy, specificity, and therapeutic potential of these compounds. Structural and mechanistic characterization of the ERM-BP compound interactions, together with subsequent lead optimization, may facilitate the development of more potent and selective inhibitors. Overall, this work provides a foundation for targeting parasite-specific transcriptional regulators as a strategy to disrupt encystation and potentially limit transmission of *Entamoeba* and the associated burden of amoebiasis.

## MATERIALS AND METHODS

### Parasite culture, transfection and stage conversion

*Entamoeba invadens* (strain IP-1) was cultured axenically under standard conditions. To facilitate encystation, trophozoites were transferred into 47% LYI-LG (low-glucose medium containing 7% adult bovine serum). (clark & diamond) Cells were adjusted to a density of 5 × 10⁵ cells/mL in this encystation medium and plated into 96-well plates. At 48 h and 72 h post-induction, cyst formation was evaluated by staining the chitin-containing cyst walls with calcofluor white, and images were captured using an ImageXpress Micro automated imaging system (Molecular Devices) at 10× magnification. Quantitative analysis of the images was carried out using MetaXpress software (Molecular Devices) (27, 28).

### Plasmid construction

To generate constructs for overexpression of wild-type ERM-BP (EIN_083100) and its mutant (ERM-BP-C198A) in *Entamoeba*, the full-length coding sequence was cloned into pGEX-4T1, a GST-fusion expression vector, using the *Bam*HI and *Not*I restriction sites. The C198A mutant was generated by substituting the cysteine residue at position 198 with alanine, using a site-directed mutagenesis kit according to the prescribed protocol. All constructs were verified by DNA sequencing prior to downstream recombinant protein expression and purification (27).

### Expression and purification of recombinant proteins

GST-tagged wild-type ERM-BP and the ERM-BP-C198A mutant protein were expressed in *Escherichia coli* BL21 cells following induction with IPTG. Cells were harvested by centrifugation, resuspended in GST binding buffer (25 mM Tris, pH 7.5, 150 mM NaCl, 10 mM MgCl₂, 5 mM DTT, 1 mM PMSF, 1× protease inhibitor cocktail [Sigma]), and lysed by sonication. Cell debris was removed by centrifugation, and the clarified lysate was incubated with pre-washed glutathione-Sepharose beads (GE Healthcare) to capture GST-tagged ERM-BP fusion proteins. The beads were washed with high-salt buffer (500 mM NaCl, 50 mM Tris-HCl, pH 7.5, 100 mM PMSF) to remove non-specifically bound proteins. Bound proteins were eluted using elution buffer (10 mM reduced glutathione, 50 mM Tris-HCl, pH 8.0, 10 mM MgCl₂, 1 mM PMSF, 1× protease inhibitor cocktail). Eluted wild-type and mutant proteins were dialyzed against dialysis buffer (5 mM HEPES, pH 7.6, 1 mM DTT, 0.2 mM PMSF, 1 mM EDTA, 10% glycerol) to remove residual glutathione. Purified proteins were quantified and their purity assessed by SDS-PAGE before use in subsequent assays (27).

### Protein thermal stability shift assay

Screening of small molecules against ERM-BP was performed using a target-based drug discovery approach utilizing Protein Thermal Stability Assay (PTSA). PTSA was performed according to the manufacturer’s instructions using the Protein Thermal Shift Starter Kit (Applied Biosystems; Cat. No. 4462263). All reactions were prepared in final volumes of 20 μl in 96-well plates. Each reaction mixture contained 1 μg purified recombinant protein (ERM-BP-WT or ERM-BP-C198A), 5 μl thermal stability buffer, and 2.5 μl of 8× fluorescent dye along with test compounds at a final concentration of 100 µM. Samples were prepared in quadruplets for each condition, and at least three biologically independent experiments were performed. The temperature gradient was programmed from 25°C to 99°C with a ramp rate of 0.05°C/s using a real-time PCR system (Applied Biosystems). Melting temperatures (Tm) were calculated using Protein Thermal Shift™ Software 1.x (27).

### Electrophoretic mobility shift assay (EMSA)

To determine whether the test compounds interfere with ERM-BP’s ability to bind its target DNA, we performed an electrophoretic mobility shift assay using purified recombinant ERM-BP protein together with P^32^-labelled DNA probe representing the ERM regulatory motif. Binding reactions were set up in EMSA-binding buffer (10 mM Tris-HCl, pH 7.9, 50 mM NaCl, 1 mM EDTA, 3% glycerol, 0.05% milk powder, and bromophenol blue), where the protein and probe were allowed to bind in the presence or absence of the test compounds PFMK, LEK, and EBLL. In the initial screen, each compound was tested at a single concentration (2 μM) alongside a control reaction containing only protein and probe, with no compound added. To look more closely at PFMK, we also set up a dose-response experiment, treating the protein-probe reaction with increasing concentrations (0, 0.1, 1, 2 and 4 μM) to see whether the effect on binding became stronger as the compound concentration increased. Once the binding reactions were complete, they were run on a native polyacrylamide gel to separate the protein-bound (shifted) probe from the unbound free probe. A weaker shifted band, along with more free probe, indicated that the compound was disrupting ERM-BP’s binding to its DNA target. (27, 28, 49)

### Cell viability and encystation efficiency assays

Cell viability of *Entamoeba invadens* trophozoites treated with the test compounds (PFMK, LEK, and EBLL) was assessed by the trypan blue exclusion assay. Trophozoites were treated with the indicated compounds across a range of concentrations (0–6 μM). After treatment, cells were harvested, then washed with 1× PBS, and stained with trypan blue dye to distinguish viable (dye-excluding) from non-viable (dye-permeable) cells. Cells were then counted using a hemocytometer, and viability was calculated using the formula: Viability (%) = (total live cells × 100) / (total live and dead cells). Viability assays were performed on three independent occasions (50)

To determine the effect of these compounds on promoting encystation, trophozoites were harvested, washed, and resuspended in low-glucose (LG) encystation media in the presence of DMSO (control), PFMK, LEK, and EBLL. Cultures were incubated under encystation-inducing conditions. Encystation efficiency was determined by treating an aliquot of the culture with sarkosyl to selectively lyse trophozoites while sparing detergent-resistant cysts, and the resulting cyst fraction was identified by light microscopy based on characteristic morphological features, including a spherical shape with a well-defined cell wall. Encystation efficiency for each treatment group was expressed as a percentage relative to the DMSO-treated control. Statistical significance between treatment groups and control was determined using an appropriate statistical test, with p < 0.05 considered significant (*p < 0.05, **p < 0.01; ns = not significant) (10)

### Staining and Fluorescence Microscopy

To determine the cyst wall morphology following treatment with PFMK, LEK, and EBLL, cysts were stained with 0.05% Calcofluor White for 10 min at room temperature followed by PI staining, with 50 µg/mL for 10–20 min at room temperature in dark. For staining with DAPI cells were fixed with 4% paraformaldehyde followed by permeabilization with 0.5% Triton X-100 before staining with 0.5 µg/ml final concentration DAPI for 5-10 min at room temperature in dark. The stained cysts were then washed with 1X PBS and were mounted on slides using Vectashield mounting medium and observed under fluorescence microscope.

### Immunostaining

After the permeabilization, the cells were blocked with 3% BSA in order to reduce non-specific binding. The localization of Jessie3 was determined by incubating the cells with an anti-Jessie3 antibody that had been conjugated to a fluorophore. The slides were mounted using a Vectashield mounting medium which included DAPI for staining the nuclei, and the cells were then examined by fluorescence microscopy. Bright-field images were obtained at the same time to allow assessment of the general cell morphology, and the images of Jessie3, DAPI and the bright-field images were combined in order to evaluate the spatial relationship between Jessie3 and the nuclear staining. In the control cysts, Jessie3 appeared as a continuous and well-defined ring around the cell periphery, this being in agreement with correct cyst wall assembly. By contrast, the cells that had been treated with PFMK and EBLL showed a disorganized and punctate distribution of Jessie3 which did not result in the formation of a continuous peripheral ring, indicating that the localization of cyst wall protein had been disrupted (28).

## ACKNOWLEDGEMENTS

We thank John Samuelson, Boston University for the kind gift of *Entamoeba* Jaccie-3 antibodies. We thank Matthew Bogyo, Stanford University for some of the Cysteine electrophile compounds and all Manna lab members for critical reading of the manuscript.

## FUNDING

This work was supported by SERB-CRG project, Govt. of India ANRF(CRG/2023/005414/BHS). S.H was supported by ANRF-JRF. S.K.D was supported from DBT-JRF, Govt. of India (DBT/2022-23/RMVERI/2110). S.P. was supported by CSIR-UGC JRF, Govt. of India.

## AUTHOR CONTRIBUTIONS

DM conceptualized and developed the Draft. S.H, S.K.D, DM performed the experiments. S.H, S.K.D, Sh.P and DM led the writing of the manuscript. All the authors S.H, S.K.D, Sh.P, D.M contributed to the content, drafting and critical review of the manuscript. S.H and D.M prepared the figures.

## ADDITIONAL FILES

**Supplemental Materials**

